# Structural basis for C-type inactivation in a Shaker family voltage gated K^+^ channel

**DOI:** 10.1101/2021.10.01.462615

**Authors:** Ravikumar Reddi, Kimberly Matulef, Erika A. Riederer, Matthew R. Whorton, Francis I. Valiyaveetil

**Author notes:** Correspondence to FIV.

## Abstract

C-type inactivation is a process by which ion flux through a voltage-gated K^+^ (K_v_) channel is regulated at the selectivity filter. While prior studies have indicated that C-type inactivation involves structural changes at the selectivity filter, the nature of the changes have not been resolved. Here we report the crystal structure of the K_v_1.2 channel in a C-type inactivated state. The structure shows that C-type inactivation involves changes in the selectivity filter that disrupt the outer two ion binding sites in the filter. The changes at the selectivity filter propagate to the extracellular mouth and the turret regions of the channel pore. The structural changes observed are consistent with the functional hallmarks of C-type inactivation. This study highlights the intricate interplay between K^+^ occupancy at the ion binding sites and the interactions of the selectivity filter in determining the balance between the conductive and the inactivated conformations of the filter.

## Introduction

Voltage gated K^+^ channels (K_v_) are essential for the generation and conduction of electrical signals by neurons, muscle and endocrine cells.(1) K_v_ channels are tetrameric proteins that contain a central pore domain and peripheral voltage sensor domains (VSD, Fig.1a).(2, 3) The pore domain houses the pathway for K^+^ ions across the membrane (Fig. 1b). The activation of a K_v_ channel takes place upon membrane depolarization and involves an outward movement of the VSDs.(4) This movement of the VSD is coupled to the opening of pore domain to turn on the K^+^ flux across the membrane. On sustained activation, the flux of K^+^ through the K_v_ channel is turned off through mechanisms that are referred to as inactivation.(5)

**Figure 1:**
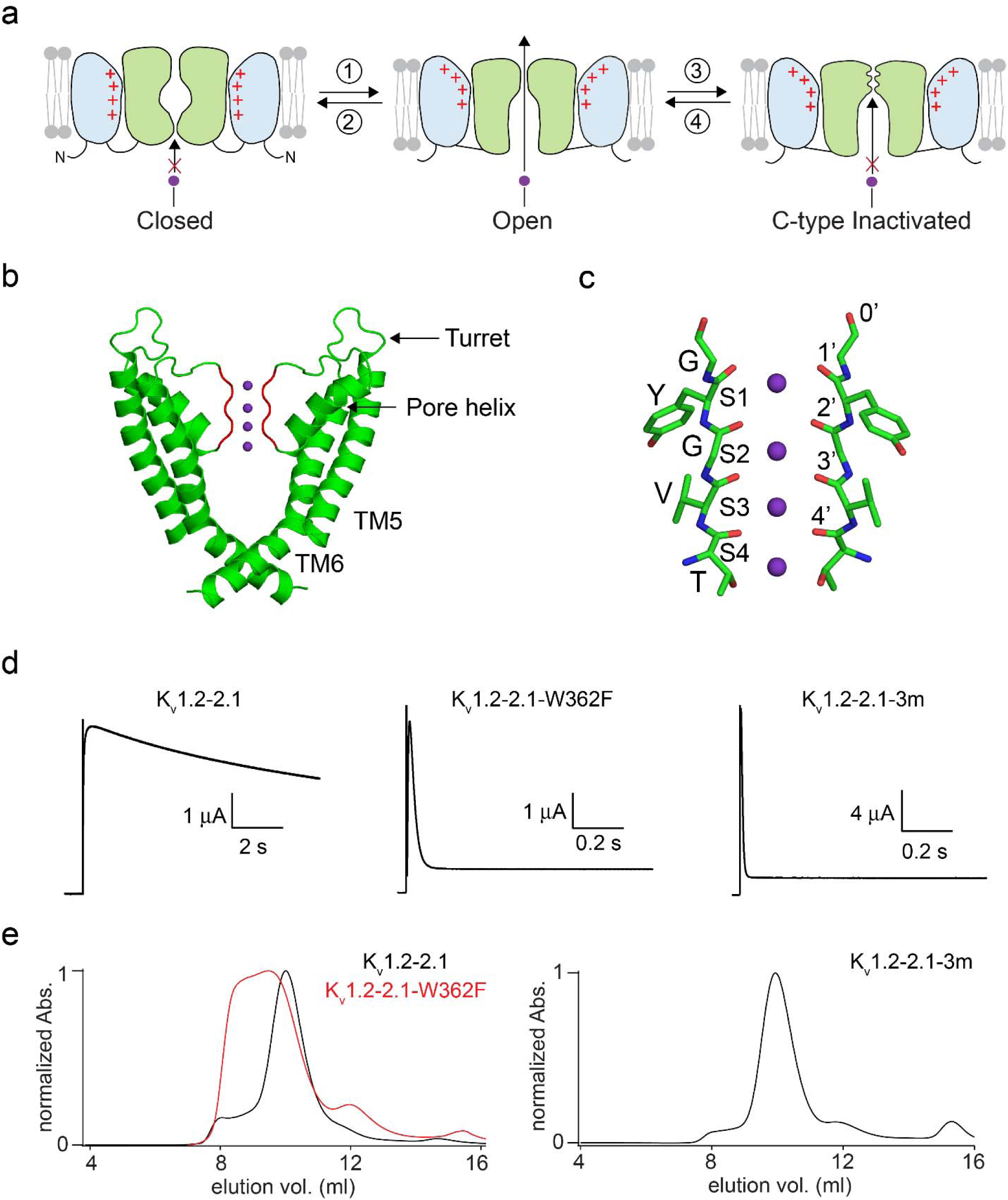
C-type inactivation in the K_v_1.2-2.1 channel. **a**) Gating mechanisms in K_v_ channels. The gating processes in a K_v_ channel of activation (1), deactivation (2), C-type inactivation (3) and recovery from inactivation (4) are depicted. TM4 in the voltage sensor domain is colored red. Voltage gating involves the movement of the TM4 helix while C-type inactivation involves structural changes at the selectivity filter. **b)** The pore domain of the K_v_1.2-2.1 channel (pdb: 2r9r). Two opposite subunits are shown, with the selectivity filter colored red and the K^+^ ions bound at the selectivity filter depicted as purple spheres. **c)** Close-up view of the selectivity filter of the K_v_1.2-2.1 channel. Two opposite subunits are shown in stick representation. The ion-binding sites in the selectivity filter (S1–S4) and the 0′–4′ carbonyl bonds are labeled. **d)** C-type inactivation in the K_v_1.2-2.1, K_v_1.2-2.1-W362F and the K_v_1.2-2.1-3m channels. Time course of current elicited by stepping the voltage from –80 mV holding potential to 40 mV with 100 mM external K^+^. Initial spikes observed are the capacitance transients. The inactivation time constants (in ms) determined were 11890 ± 1666 (± SEM, n = 4) for K_v_1.2-2.1, 22.4 ± 7.2 (9) for the K_v_1.2-2.1-W362F and 5.5 ± 0.7 (10) for the K_v_1.2-2.1-3m channels. **e**) Size exclusion chromatography profiles of the purified K_v_1.2-2.1, K_v_1.2-2.1-W362F and the K_v_1.2-2.1-3m channel.

In the Shaker family of K_v_ channels, there are two types of inactivation, N-type and C-type.(6, 7) In N-type inactivation, the N-terminus of the channel binds to the open pore domain and occludes the ion pathway.(6, 8) Shaker channels with the N-terminal deleted do not undergo N-type inactivation but inactivate through a different mechanism called C-type inactivation.(7) This mechanism is also referred to as slow inactivation as the timescale is generally slower, on the order of seconds, compared to the millisecond time scale observed for N-type inactivation.(9) C-type inactivation is a physiologically important process that can regulate cell excitability by determining channel availability.(10, 11)

The mechanism of C-type inactivation has been investigated using functional, spectroscopic, structural and computational approaches.(5) All these approaches suggest that C-type inactivation involves changes at the selectivity filter. The selectivity filter refers to the narrow region of the ion pathway in the pore domain, where selection for K^+^ takes place (Fig. 1c).(12) The selectivity filter consists of a row of K^+^ binding sites (called S1-S4 extracellular – intracellular) that are constructed from the main chain carbonyl atoms and the threonine side chain from the protein sequence T-V-G-Y-G.(13) The structures of the selectivity filter of K^+^ channels are highly conserved. The structures determined mainly show the selectivity filter in a conductive state, the state of the selectivity filter that supports the flux of K^+^ through the channel. During C-type inactivation, there are structural changes at the selectivity filter that convert the selectivity filter from a conductive to a non-conductive conformation.(5, 9)

Understanding the mechanism of C-type inactivation requires the structure of the selectivity filter in the C-type inactivated state. The structure of a non-conducting mutant of the K_v_1.2 channel has been proposed to show the selectivity filter in the C-type inactivated state.(14) However, the structural changes observed were minimal, which raises a question on whether this structure truly represents the C-type inactivated state.(15) Other structural studies on C-type inactivation have used the K^+^ channel KcsA. KcsA does not belong to the K_v_ family but undergoes an inactivation process that bears some functional similarity to C-type inactivation in K_v_ channels.(16) Structural studies on the KcsA channel under conditions that favor inactivation show a constriction of the selectivity filter.(17, 18) However, the relevance of this constricted conformation to C-type inactivation in K_v_ channels has been debated.(19)

Here we report the crystal structure of the K_v_1.2 channel in the C-type inactivated state. We carry out structural studies on a mutant of the K_v_1.2 channel that shows a vastly increased rate of C-type inactivation. We observe that C-type inactivation involves a dilation of the outer ion binding sites of the selectivity filter along with changes in the extracellular mouth and the turret regions of the pore domain. Our studies highlight the selectivity filter interactions that are important for C-type inactivation and suggest a molecular pathway for this process.

## Results

### C-type inactivation in the K_v_1.2-2.1 channel

Our strategy for determining the structure of the K_v_1.2 channel in the C-type inactivated state was to use mutants of this channel that show an enhanced rate of inactivation with the expectation that these mutants will trap the selectivity filter in the C-type inactivated state. In the Shaker channel, the W434F substitution dramatically increases the rate of inactivation and this substitution has been extensively used in functional studies to mimic the C-type inactivated state.(20) The corresponding mutation, W366F in the K_v_1.2 channel has been shown to have an increased rate of inactivation (Supplementary fig. 1).(21, 22) For our studies, we used a variant of the K_v_1.2 channel referred to as the K_v_1.2-2.1 chimera (referred henceforth as K_v_1.2-2.1) as this construct affords crystals that diffract to a higher resolution.(3) In the K_v_1.2-2.1 channel, residues 267-302 in the S3-S4 loop in the voltage sensor domain are substituted by residues 274-305 of the K_v_2.1 channel. The K_v_1.2-2.1 channel shows similar functional properties to the K_v_1.2 channel and the W362F (equivalent to W366F in K_v_1.2) substitution in the K_v_1.2-2.1 channel shows a similar enhancement in the rate of inactivation (Fig. 1d).(23)

Our attempts at structural studies of the K_v_1.2-2.1-W362F channel were stymied as the channel showed poor expression and suboptimal biochemistry (Fig. 1e). We therefore screened for additional amino acid substitutions that improved protein expression and biochemical behavior while maintaining an enhanced rate of C-type inactivation. These efforts identified a construct with two additional substitutions, S367T and V377T, which exhibited good biochemical behavior and an enhanced rate of inactivation (Fig. 1d, e). We refer to the K_v_1.2-2.1 channel with the W362F, S367T and V377T substitutions as the K_v_1.2-2.1-3m channel. At high K^+^, the K_v_1.2-2.1-3m channel showed a ~2000-fold increase in the rate of C-type inactivation while at low external K^+^, the rate of C-type inactivation was very fast and could not be accurately determined. Based on the rapid rate of inactivation, we anticipated that the crystal structure of the K_v_1.2-2.1-3m channel would reveal the selectivity filter in the C-type inactivated state.

### Structure of the K_v_1.2-2.1-3m channel

For structural studies, the K_v_1.2-2.1-3m channel was co-expressed with the beta subunit and the channel-beta complex was crystallized in the presence of 150 mM K^+^. We obtained crystals of the same space group (*P4 2*_*1*_ *2*) as reported for the K_v_1.2-2.1 channel but with slightly altered unit cell parameters.(3) Data from multiple crystals were combined to obtain a complete data set to 3.35 Å resolution and the structure was determined by molecular replacement using the K_v_1.2-2.1 (pdb: 2r9r) channel structure as the model (Supplementary table 1). There are two copies (molecule I and II) of the channel-beta complex in the asymmetric unit. The beta subunit and the T1 domain in both molecules were well resolved while the transmembrane region in molecule I was better resolved than in molecule II. For the transmembrane region, we observed stronger electron density for the pore domain compared to the voltage sensor. Within the pore domain, the electron density for Y373-D375 was weaker compared to the neighboring residues and electron density for the carbonyl group for the G374 residue was not observed. Electron density corresponding to the pore helix and the selectivity filter are shown in figure 2a, b. In the voltage sensor domain, the electron density was sufficient for modelling the transmembrane segments (Supplementary fig. 2) while electron density for the loops connecting the TM segments and the linker connecting the T1 domain to the first transmembrane segment were not observed and these regions were therefore not modelled. A superposition of the various domains in the K_v_1.2-2.1-3m channel to K_v_1.2-2.1 channel shows that the structural changes observed are mainly in the selectivity filter region and in the extracellular mouth of the pore (Supplementary fig. 3).

**Figure 2:**
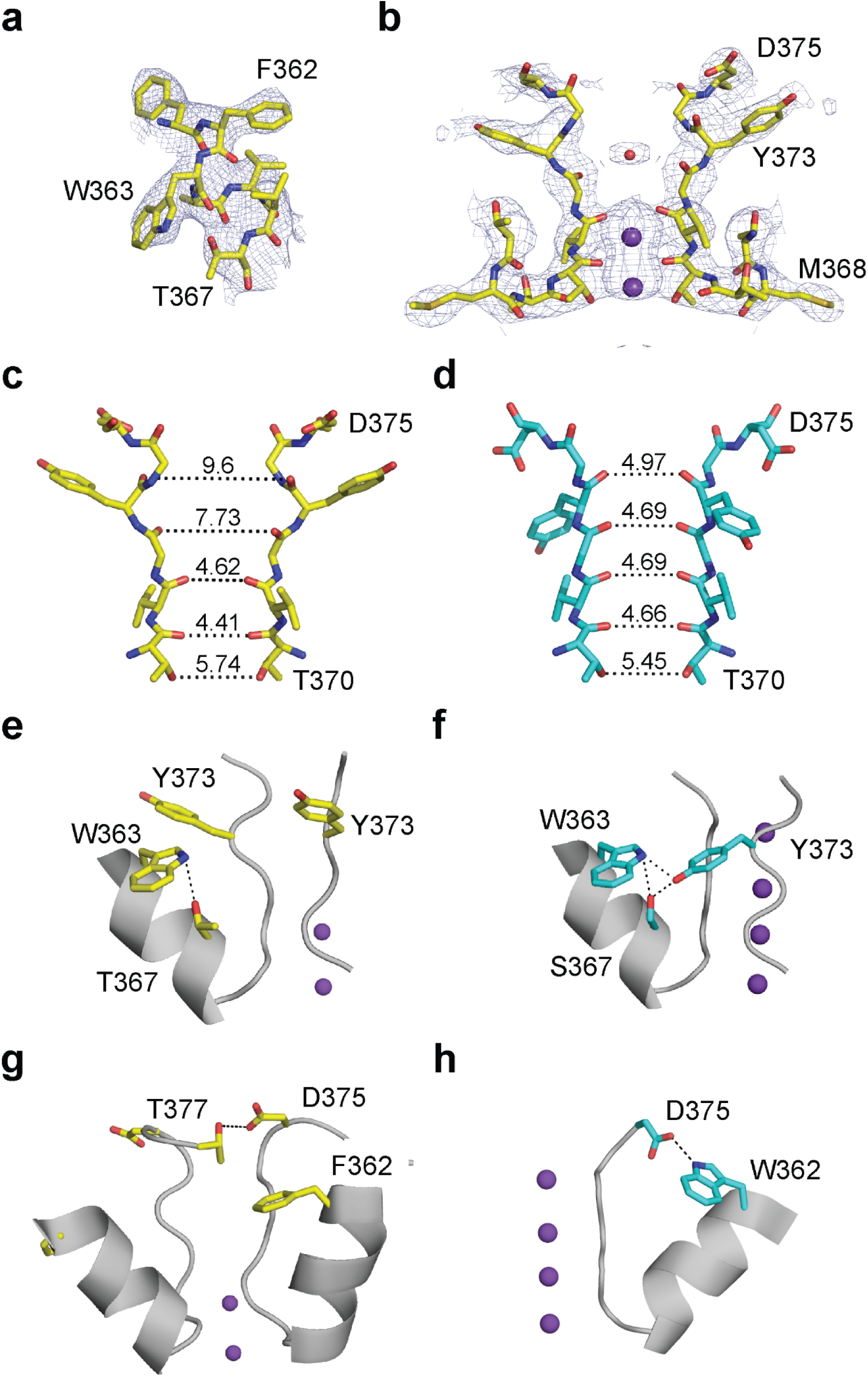
Changes in the selectivity filter on C-type inactivation. **a**) The pore helix of the K_v_1.2-2.1-3m channel. The 2F_o_-F_c_ electron density map contoured at 1.4 σ is shown with residues 361-367 as sticks. The W362F and S367T substitutions present in the K_v_1.2-2.1-3m channel are indicated. **b**) The selectivity filter of the K_v_1.2-2.1-3m channel. The 2F_o_-F_c_ electron density map contoured at 0.9 σ is shown with residues 368-375 as sticks, K^+^ ions as purple spheres and water molecule as a red sphere. **c**,**d)** Comparison of the selectivity filter of the K_v_1.2-2.1-3m (**c**) and the K_v_1.2-2.1 channel (**d**). Residues 370-375 are shown as sticks. The distance (in Å) between the carbonyl oxygens and the Thr370 side chain hydroxyl group in the opposite subunits are indicated by dotted lines. **e**,**f**,**g**,**h**) Close-up of the selectivity filter and the pore helix showing the Y373 side chain in the K_v_1.2-2.1-3m (**e**) and K_v_1.2-2.1 channel (**f**) and the D375 side chain in the K_v_1.2-2.1-3m (**g**) and K_v_1.2-2.1 channel (**h**).

### The selectivity filter in the K_v_1.2-2.1-3m channel

A comparison of the selectivity filter in K_v_1.2-2.1-3m to the wild type channel shows substantial changes in the sidechain and backbone conformations (Fig. 2c, d). A major change observed is a reorientation of the Tyr side chain (Fig. 2e, f). In the K_v_1.2-2.1 channel, the Tyr side chain interacts with the Trp363 and Ser 367 of the adjacent subunit while in the K_v_1.2-2.1-3m channel, these H-bond interactions are broken and the Tyr side chain undergoes a 77° rotation and the hydroxyl group in the Tyr side chain is now oriented towards the extracellular surface.

Another major change observed in the K_v_1.2-2.1-3m channel is a flip of the Asp375 side chain (Fig. 2g, h). In the K_v_1.2-2.1 structure, Asp375 forms an H-bond with Trp362 in the pore helix. This H-bond is disrupted in the K_v_1.2-2.1-3m channel due to the W362F substitution. The Asp375 side chain is now oriented towards the extracellular solution and forms a H-bond with Thr377 in the adjacent subunit (distance of 3.32 Å). The K_v_1.2-2.1 channel has a Val at this position, which is substituted by Thr in the K_v_1.2-2.1-3m channel. The flip of the Asp side chain as observed changes the side chain from being oriented towards the channel interior (in K_v_1.2-2.1) to facing the extracellular solution.

The changes in the Tyr373 and Asp375 side chain conformations cause a widening of the selectivity filter towards the extracellular side. Due to this widening, the S1 and S2 ion binding sites are disrupted and instead form a vestibule (Fig. 2c, d). In K_v_1.2-2.1, K^+^ binding is observed at the S1 and the S2 sites while in the K_v_1.2-2.1-3m channel only a weak electron density is observed in this vestibule region (Fig. 2b, Supplementary fig. 4). The electron density in vestibule region can correspond either to a low occupancy K^+^ ion or a water molecule. The distances between the carbonyl oxygens in the vestibule region are however too large to tightly coordinate either a water or a K^+^ ion. In contrast to the changes seen at the S1 and the S2 sites, the S3 and the S4 sites in the K_v_1.2-2.1-3m filter are well superimposed on the corresponding sites in the K_v_1.2-2.1 filter and electron density corresponding to ion binding at these sites is observed. Previous studies have implicated the outer sites in the selectivity filter in C-type inactivation (24, 25) and the K_v_1.2-2.1-3m structure indicates that inactivation involves a disruption of the S1 and the S2 sites in the selectivity filter. A key characteristic of C-type inactivation is a dependence on the extracellular K^+^ concentration.(26, 27) The structure suggests that extracellular K^+^ is protective for the filter against C-type inactivation as higher extracellular K^+^ concentration will increase the ion occupancy at the S1/S2 sites to prevent the dilation of these sites.

### Changes in the extracellular mouth of the K_v_1.2-2.1-3m pore

The changes in the selectivity filter in the K_v_1.2-2.1-3m structure are propagated to the loop region linking the selectivity filter to TM6 (Fig. 3a). The changes observed in this region are mainly due to the repositioning of the Asp375 side chain (Fig. 3b,c). Studies on the Shaker K^+^ channel have suggested that residues corresponding to 376-378 in the K_v_1.2-2.1-3m channel change accessibility on C-type inactivation.(28) Studies on Shaker have also shown that a Cys substitution at 448 (376 in K_v_1.2-2.1-3m) exhibits an enhanced rate of formation of a disulfide bond while a Cys substitution at 449 (377 in K_v_1.2-2.1-3m) can form a high affinity metal (Cd^2+^ or Zn^2+^) binding site in the inactivated state.(28, 29) Comparison of the 376-378 region in the K_v_1.2-2.1-3m to the K_v_1.2-2.1 shows an increase in surface exposure of these residues (Supplementary fig. 5), which may underlie the changes in modification observed (for Cys substitutions at these sites) on C-type inactivation. In the K_v_1.2-2.1-3m structure, the Cα-Cα distance between the 376 sidechains in the adjacent subunits is similar to the distance observed in the K_v_1.2-2.1 structure (Fig. 3b, c). The K_v_1.2-2.1-3m structure therefore does not account for the enhanced rate of disulfide bond formation by Cys 376 in the C-type inactivated state. Similarly the change in the Cα-Cα distance of the 377 sidechains in these structures is 0.51 Å (14.06 to 13.55 Å), and this distance is still longer than expected for the formation of a metal coordination site by the Cys 377 sidechains from adjacent subunits.(30) The K_v_1.2-2.1-3m channel structure therefore does not directly account for some of the changes in the extracellular mouth of the pore on C-type inactivation as inferred from the functional studies. One possibility is that the full extent of changes in the extracellular mouth of the pore on C-type inactivation are not revealed in the K_v_1.2-2.1-3m structure. Another possibility is that these experimental findings reflect greater flexibility in the extracellular mouth of the channel in the C-type inactivated state. An increased flexibility in the extracellular mouth on C-type inactivation could explain the poor electron density observed in the Y373-D375 region in the K_v_1.2-2.1-3m channel.

**Figure 3:**
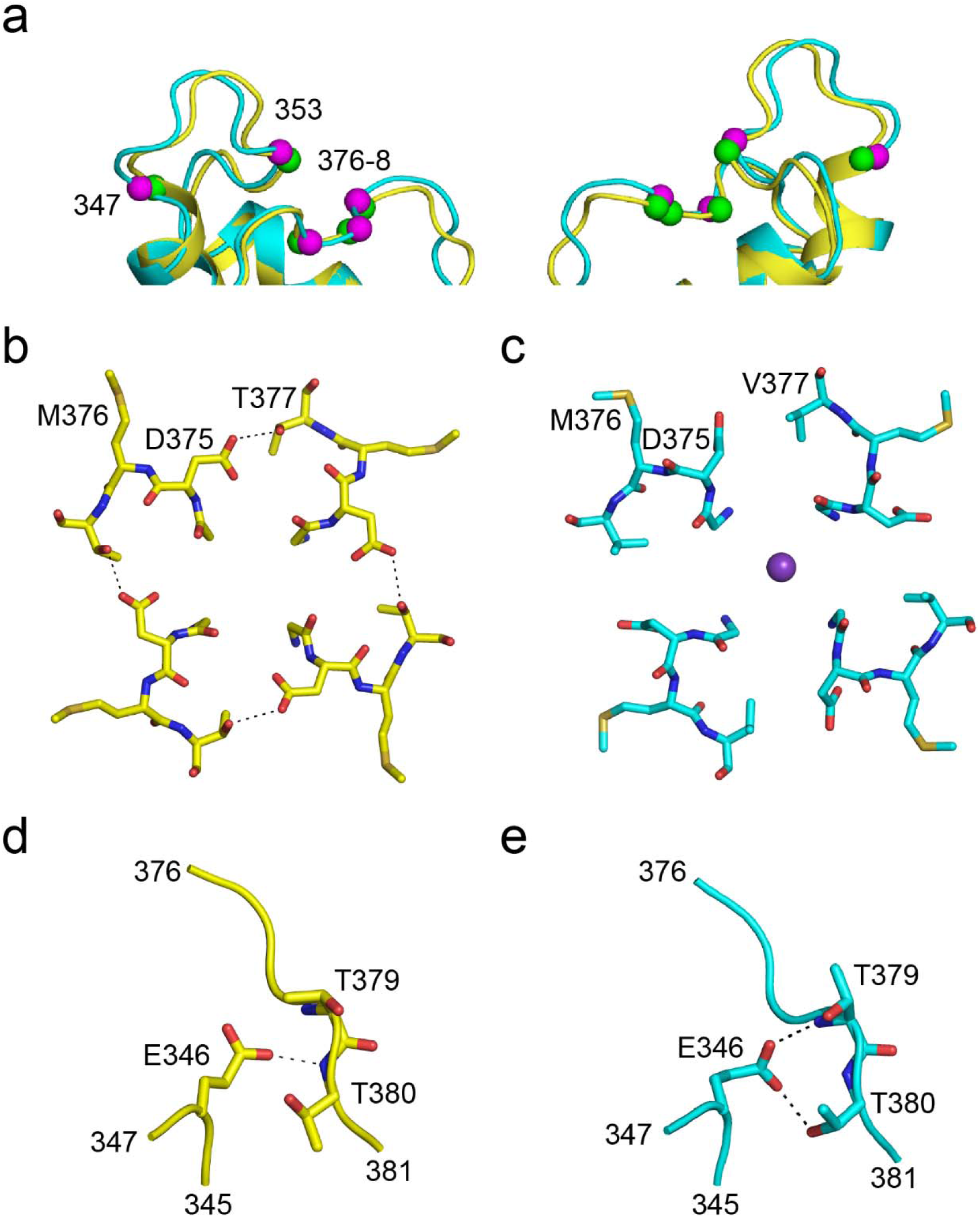
The outer pore region in the C-type inactivated channel. **a**) Superposition of the outer pore region of the K_v_2.1-2.1-3m (yellow) and the K_v_2.1-2.1 channel (cyan). Two opposite subunits are shown. Some of the positions at which fluorescence probes introduced detect structural changes on C-type inactivation are indicated (green spheres in K_v_2.1-2.1-3m and magenta spheres in K_v_2.1-2.1). **b**,**c)** Top view of the extracellular mouth of the pore in the K_v_2.1-2.1-3m (**b**) and the K_v_1.2-2.1 channel (**c**) shows changes in conformation of D375 and M376. The H-bond interaction between D375 and T377 in adjacent subunits of the K_v_2.1-2.1-3m channel is shown. **d**,**e**) H-bond interactions of E346 in K_v_1.2-2.1-3m (**d**) and K_v_1.2-2.1 channel (**e**) are shown.

C-type inactivation also involves changes in the turret region present between TM5 and the pore helix. Fluorescent probes introduced into the turret region show a change in fluorescence on C-type inactivation.(31–33) We see structural changes in the turret region of the K_v_1.2-2.1-3m corresponding to the sites of introduction of the fluorescence probes (Fig. 3a). A key interaction that is altered in the K_v_1.2-2.1-3m channel is that of the E346 residue, which is towards the top of TM5 (Fig. 3d, e). Mutations at this Glu (E418 in the Shaker channel) increase the rate of inactivation.(33, 34) In the K_v_1.2-2.1 channel, E346 interacts with the pore-S6 loop region of the channel through H-bond interactions with the protein backbone of Thr379 and the side chain of Thr380. In the K_v_1.2-2.1-3m structure, we observe a change in the E346 side chain conformation and correspondingly the interactions of the E346 side chain with the T379-T380 residues are altered.

### Structure of the K_v_1.2-2.1-W362F, S367T channel

To test the importance of the interaction between Asp375 and Thr377 for C-type inactivation, we investigated the functional properties of a K_v_1.2-2.1 channel with the native Val at 377 along with the W362F and S367T substitution (referred to as K_v_1.2-2.1-2m). We observed that the inactivation was slow in this mutant compared to the very rapid inactivation observed in the K_v_1.2-2.1-3m channel. Inactivation in the K_v_1.2-2.1-2m channel was similar to the inactivation observed in the Kv1.2-2.1 channel (Fig. 4a). We crystallized and determined the structure of the K_v_1.2-2.1-2m channel at 150 mM K^+^ (Fig. 4b, Supplementary table 1). In the K_v_1.2-2.1-2m structure, we observed the selectivity filter in a conductive conformation as anticipated based on the similarity of the inactivation properties of the K_v_1.2-2.1-2m to the K_v_1.2-2.1 channel (Fig. 4c). These findings suggest a role for the inter-subunit Asp375-Thr377 interaction in stabilizing the selectivity filter in the C-type inactivated state. The equivalent position in the Shaker channel (Thr449) is an important locus for C-type inactivation and substitution of T449 in the Shaker channel with Val results in a non-inactivating phenotype.(26) Our studies suggests that substitutions at this site affect C-type inactivation through an effect on the interaction with the Asp residue.

**Figure 4:**
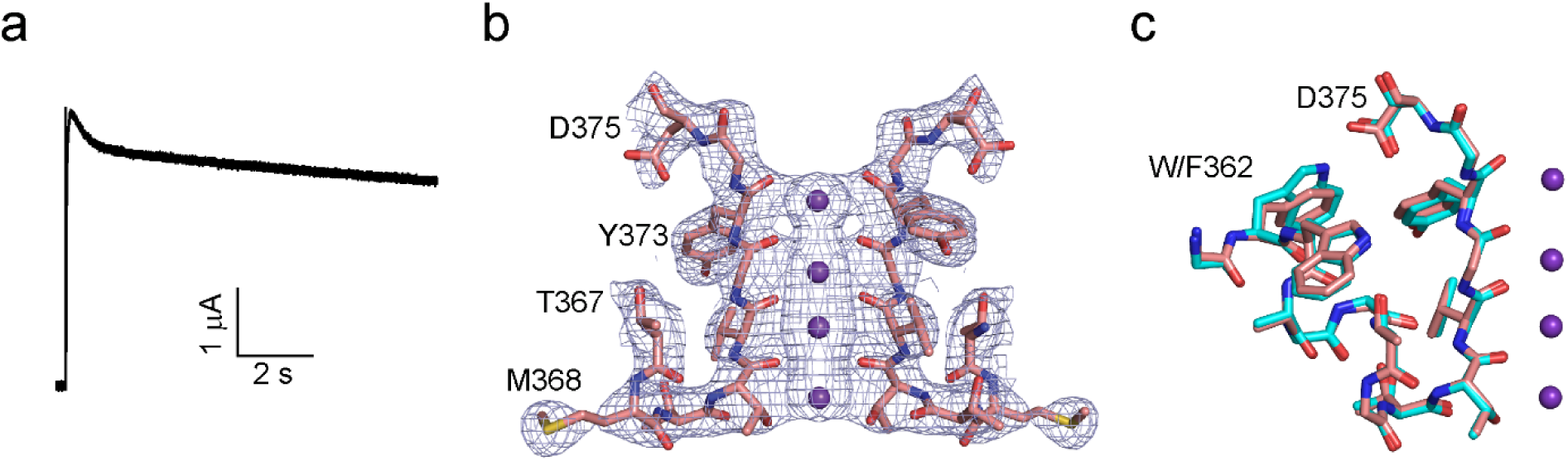
Structure of the selectivity filter in the K_v_2.1ch-2m channel. **a)** C-type inactivation in the K_v_1.2-2.1-2m (W362F, S367T) channel. Time course of the current elicited by stepping the voltage from –80 mV holding potential to 40 mV in 100 mM external K^+^. **b**) Electron density of the selectivity filter of K_v_1.2-2.1-2m. The 2F_o_-F_c_ electron density map contoured at 1.3 σ is shown with residues 367-375 as sticks, and the K^+^ ions in the selectivity filter shown as purple spheres. **c**) Superposition of the pore helix and filter region (residues 362-375) of the K_v_1.2-2.1-2m and the K_v_1.2-2.1 channel.

## Discussion

Here we report the structure of the K_v_1.2 channel in the C-type inactivated state (Fig. 5a). The structure shows that the C-type inactivation involves a dilation of the selectivity filter at the S1 and S2 ion binding sites, which is caused by structural changes at the conserved Tyr and Asp side chains. The disruption of the S1 and the S2 ion binding sites of the filter perturbs K^+^ flux through the channel. The structure shows that the changes in the selectivity filter on C-type inactivation are propagated to the extracellular mouth and the turret region of the pore domain of the channel. The structure of the selectivity filter in the C-type inactivated state is consistent with the dilation model for inactivation previously proposed by Hoshi and Armstrong.(9)

**Figure 5:**
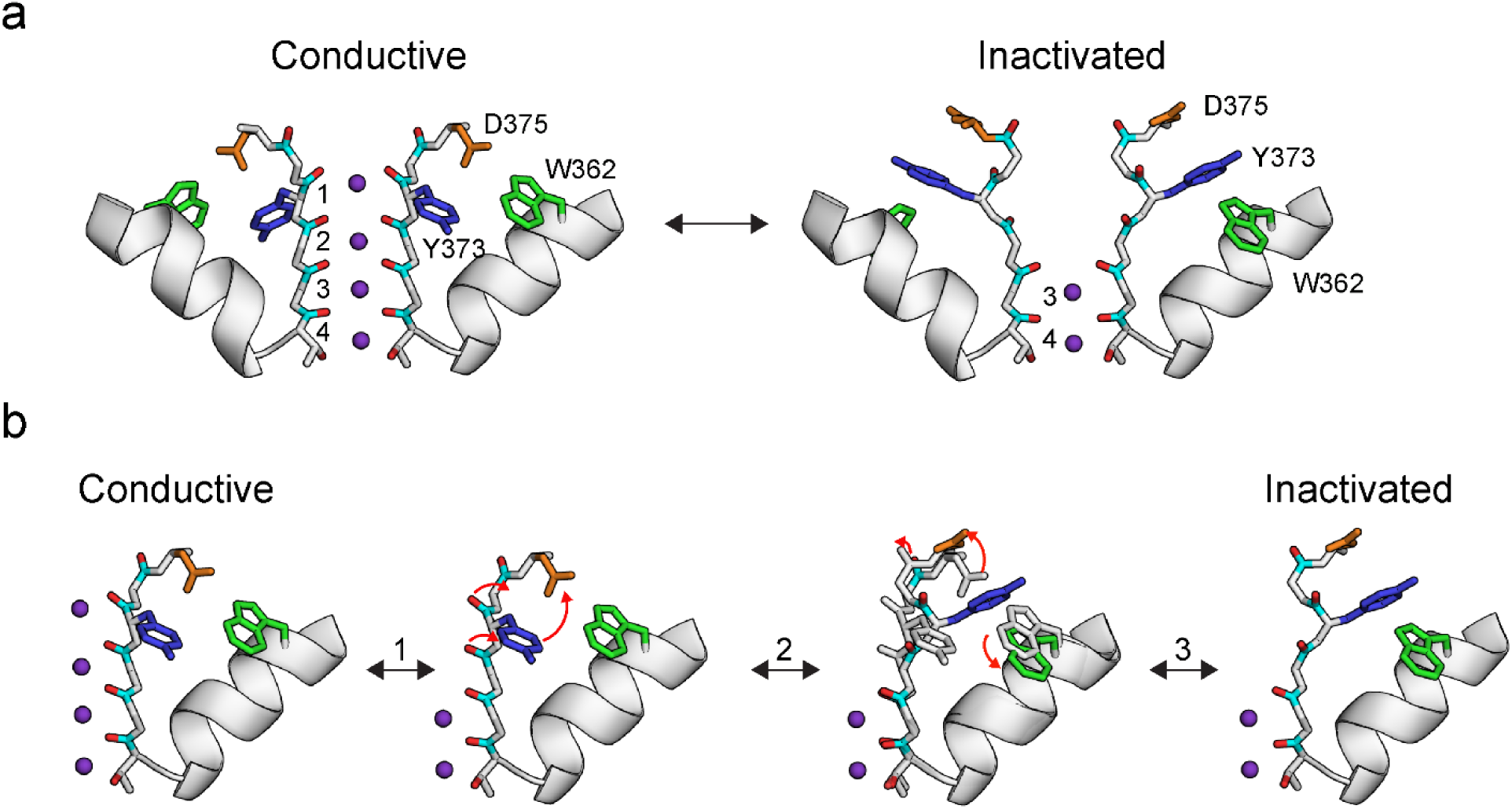
C-type inactivation in a K_v_ channel. **a)** A model for the selectivity filter in the conductive and the C-type inactivated state. The conductive state of the selectivity filter is observed in the K_v_1.2-2.1 channel (pdb: 2r9r) while the C-type inactivated state is based on the structure of the selectivity filter in the K_v_1.2-2.1-3m channel. Two opposite subunits are shown **b**) A potential sequence of structural transitions in the selectivity filter during C-type inactivation. Step 1 is the loss of ion binding to the S1 and S2 sites. Step 2 is the switch of the Y373 residue and the flip of the 1’ and the 2’ carbonyl groups. Step 3 is the rotation of the D375 residue to the extracellular side and the flip of the 0’ carbonyl group. These transitions result in a dilation of the S1 and S2 ion binding sites in the selectivity filter. One subunit is shown.

Based on our structural studies, we propose the following transitions at the selectivity filter during C-type inactivation (Fig. 5b). The process initiates with a loss of ion occupancy at the S1 and S2 sites in the selectivity filter. The loss of ion occupancy causes a flip of 1’ and the 2’ carbonyl groups, which is accompanied by a rotation of the Tyr373 side chain. The movement of the Tyr side chain sets up a clash with the Asp375 sidechain and as a result, the Asp375-Trp362 H bond interaction is broken and the Asp side chain switches to facing the extracellular side. Asp375 in this conformation is stabilized by an interaction with Thr377. This change in the conformation of Asp375 flips the 0’ carbonyl group. The changes in the 0’ – 2’ carbonyl groups result in a dilation of the S1 and the S2 sites in the selectivity filter, which prevents ion coordination at these sites and thereby affects ion conduction through the channel. The changes in the Tyr and the Asp side chain also change the surface accessibility and dynamics in the filter-TM6 region. These changes are also transmitted to the pore turret through changes in the interaction of E346 with the filter-TM6 loop. Through this process, structural changes that are initiated by changes in ion occupancy at the selectivity filter are propagated throughout the selectivity filter and the extracellular mouth of the pore.

A structure of the K_v_1.2-2.1-V406W channel has been reported as presenting the selectivity filter in the C-type inactivated state (Supplementary fig. 6).(14) The structural change observed in the selectivity filter, compared to the conductive state, is a small distortion of the S1 site.(15) This is in contrast to the abrogation of the S1 and S2 sites observed in the K_v_1.2-2.1-3m structure. The proposed inactivated state in the KcsA channel shows a distortion of the S2 and the S3 sites of the selectivity filter and is distinct from the structure of the selectivity filter in the K_v_1.2-2.1-3m channel (Supplementary fig. 6). Gating of K^+^ flux through two-pore K^+^ (K2P) channels takes place at the selectivity filter by a mechanism that shares a functional resemblance to C-type inactivation in K_v_ channels.(35) A recent structural study on the TREK-1 K2P channel suggested that the gating process involves a loss of the S1 and the S2 ion binding sites in the selectivity filter while a structure of the TASK 2-P K^+^ channel indicated a loss of the S1 (and potentially the S0) binding site in the inactivated state.(36, 37) These structures suggest that K^+^ channels, which have essentially identical selectivity filters, can have distinct mechanisms of inactivation or gating at the selectivity filter.

C-type inactivation is widespread in the K_v_ channel family with the inactivation properties that are tuned for the specific physiological roles of these channels. Structural studies of these other K_v_ family members will be necessary to determine whether conformational changes similar to that observed in the K_v_1.2-2.1-3m structure underlie inactivation or gating at the selectivity filter in diverse K_v_ channels.

## Acknowledgements

We thank Dr. Roderick Mackinnon for providing the K_v_1.2-2.1/beta plasmid. We thank Dr. Shivani Ahuja and Dr. Kim Hartfield for help with protein expression in *Pichia*, protein purification and analysis. We thank Dr. Eric Gouaux for providing access to crystallization equipment. Crystallography data were collected at GM/CA beamlines 23ID-B and 23ID-D at the Advanced Photon Source at Argonne National Laboratory and we thank the staff at the beamlines for their support with data collection. GM/CA @ APS has been funded in whole or in part with Federal funds from the National Cancer Institute (Y1-CO-1020) and the National Institute of General Medical Science (Y1-GM-1104). Use of the Advanced Photon Source was supported by the U.S. Department of Energy, Basic Energy Sciences, Office of Science, under contract No. W-31-109-ENG-38. This research was supported by a grant from the National Institute of General Medical Sciences (R01GM087546) of the National Institutes of Health to FIV. EAR was supported by a pre-doctoral fellowship from the American Heart Association (AHA 19PRE34380950).

