## Supplementary Materials for "Structural basis for C-type inactivation in a Shaker family voltage gated K^+^ channel"

**Contents**:

1) Supplementary figures 1-6

2) Supplementary Table 1

3) Methods

**Supplementary Figures**

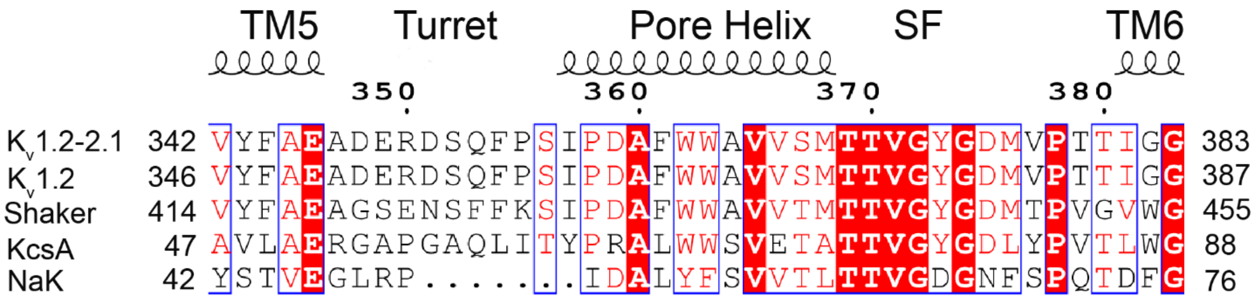


**Supplementary Figure 1**: **Sequence Alignment**. Alignment of the turret, pore helix and the selectivity filter (SF) of the $K_v$1.2-2.1 (gi: 160877794), $K_v$1.2 (gi:1235594), Shaker (gi: 13432103), KcsA (gi: 61226909) and NaK (gi: 90108700) channels. Residue numbering corresponds to the $K_v$1.2-2.1 channel. Conserved residues are highlighted in red.

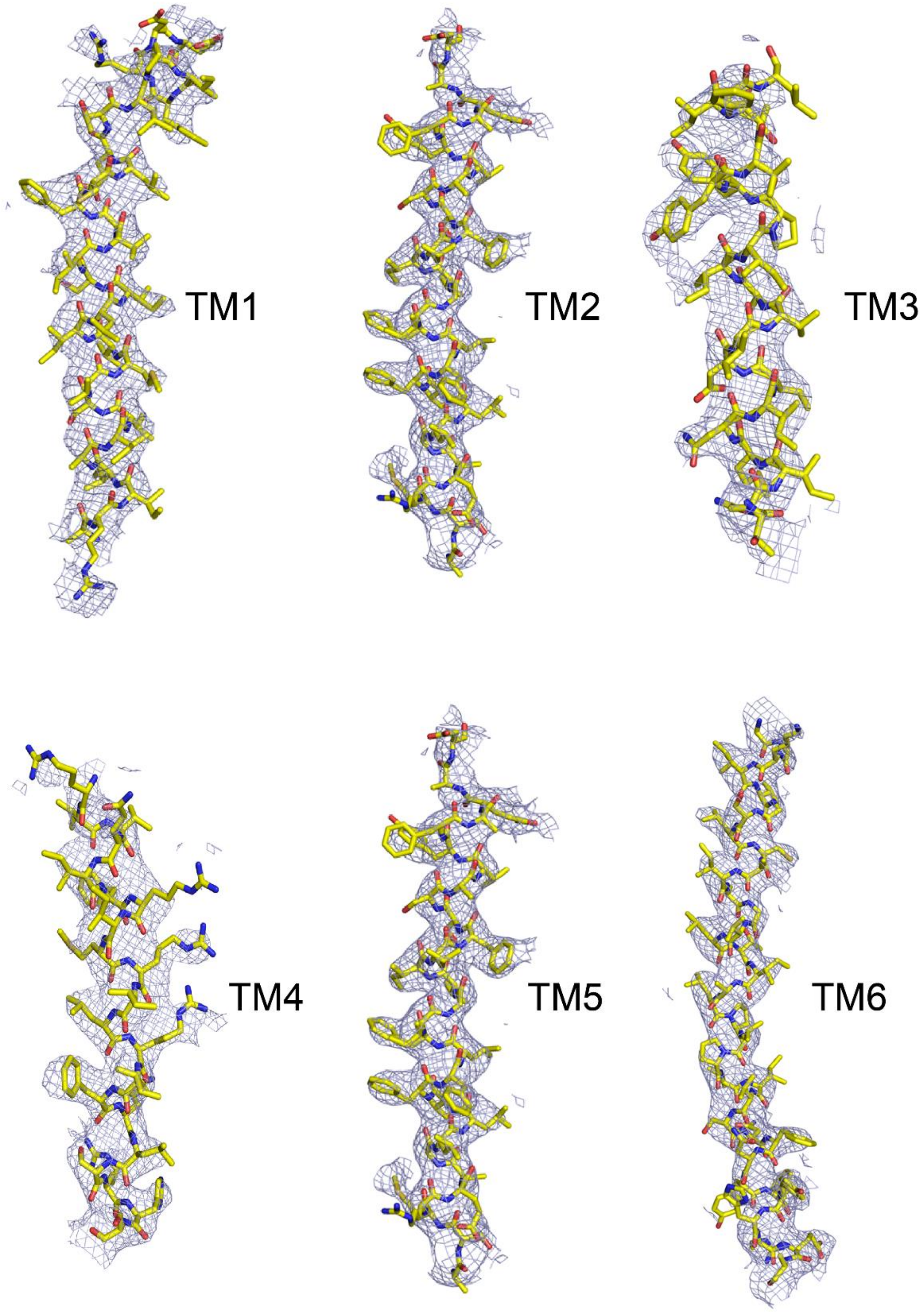


**Supplementary Figure 2**: **Transmembrane segments in the $K_v$1.2-2.1-3m channel**. 2$F_o$-$F_c$ electron density map contoured at 1 σ with the $K_v$1.2-2.1-3m structural model shown in stick representation.

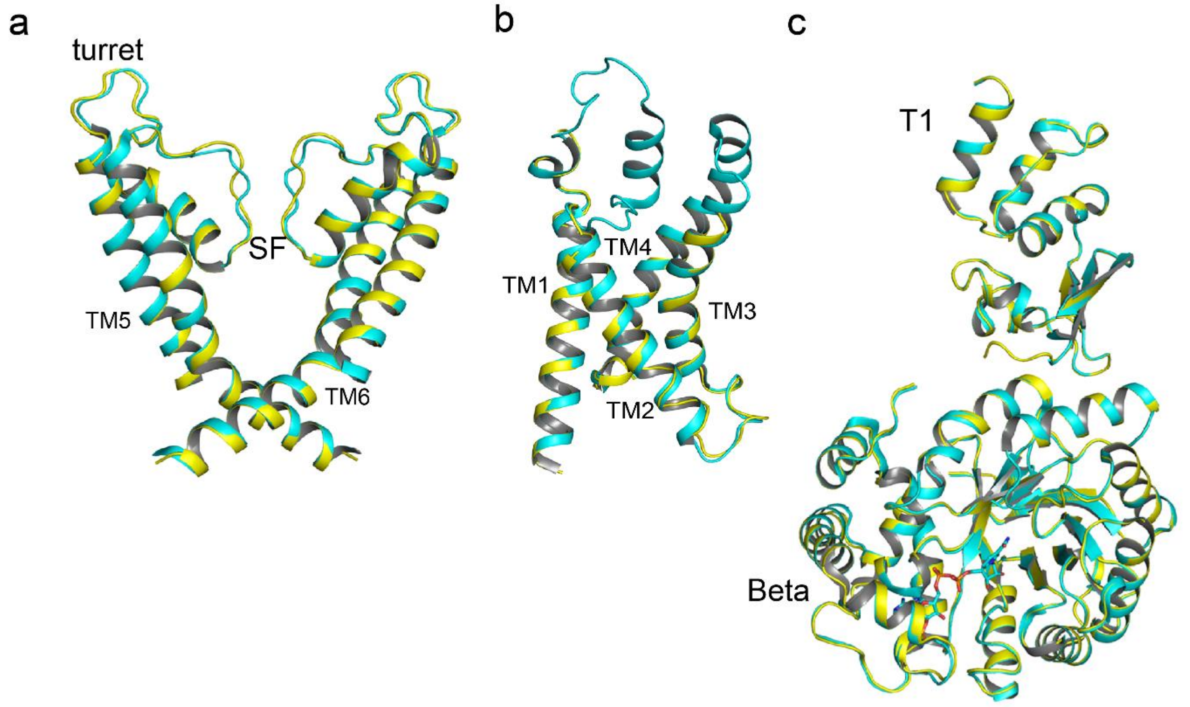


**Supplementary Figure 3**. **Comparison of $K_v$1.2-2.1-3m to $K_v$1.2-2.1.** Superposition of the pore region (**a**), the voltage sensor domain (**b**) and the T1 and beta subunits (**c**) in the $K_v$1.2-2.1-3m (yellow) and the $K_v$1.2-2.1 channel ( pdb: 2r9r, cyan) are shown. Only two opposite subunits are shown in **a**.

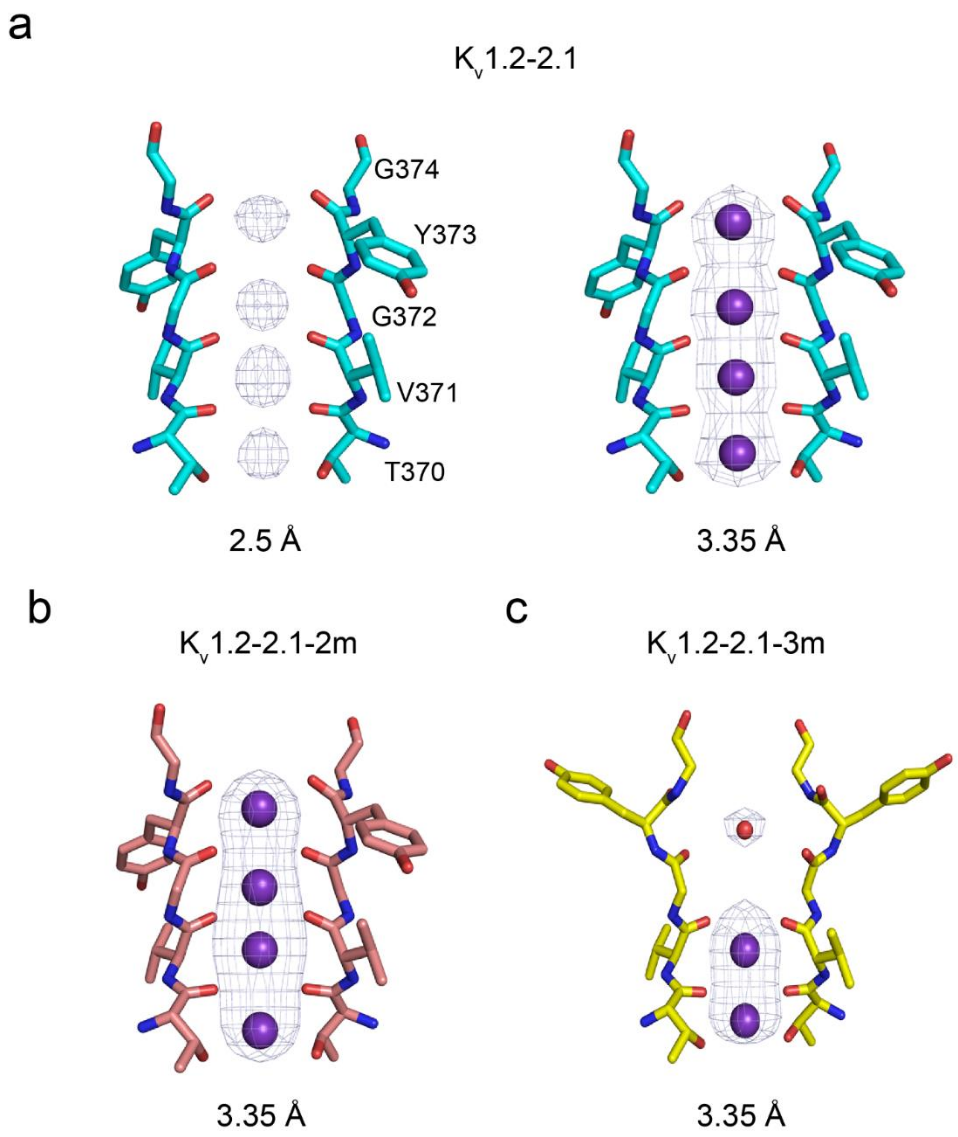


**Supplementary Figure 4. Ion binding to the selectivity filters of the $K_v$1.2-2.1, $K_v$1.2-2.1-2m and the $K_v$1.2-2.1-3m channels**. The $F_o$-$F_c$ electron density (ions were omitted) contoured at 5 $\sigma$ along the central axis of the selectivity filter is shown. Electron density calculated using diffraction data to 2.5 Å (**a,** left) and to 3.35 Å (**a,** right) for the $K_v$1.2-2.1 channel are shown. Diffraction data to 3.35 Å was used for electron density calculation for $K_v$1.2-2.1-2m (**b**) and $K_v$1.2-2.1-3m **(c)**.

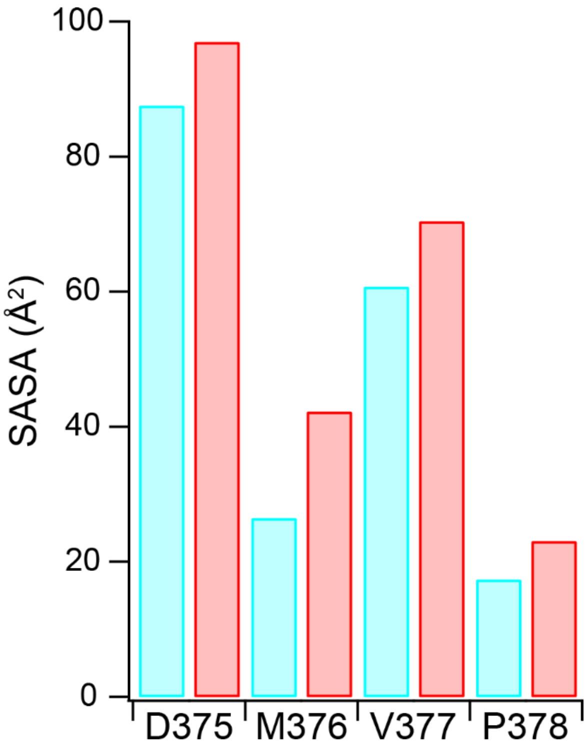


**Supplementary Figure 5**: **Changes in solvent exposure at the extracellular mouth of the pore.** Solvent accessible surface area (SASA) for residues D375-P378 at the extracellular mouth of the pore in the $K_v$1.2-2.1 (cyan) and the $K_v$1.2-2.1-3m (red) channels is shown.

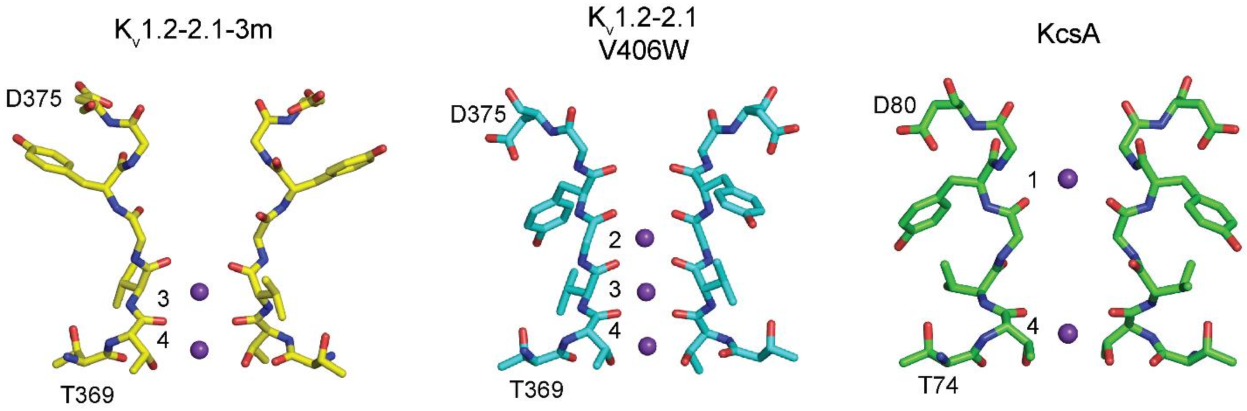


**Supplementary Figure 6**: **Gating at the K⁺ selectivity filter**. Structure of the selectivity filter of the $K_v$1.2-2.1-3m, $K_v$1.2-2.1-V406W (pdb:5wie), and the KcsA channels (pdb:5vke). Two opposite subunits are shown with the $K^+$ ions indicated by purple spheres and the binding sites are numbered.

**Supplementary Table 1**: Data collection and Crystallography statistics.

| | $K_v$1.2-2.1-3m | $K_v$1.2-2.1-2m |
|---|---|---|
| Space group | $P42_12$ | $P42_12$ |
| *a* (Å) | 129.59 | 130.7 |
| *b* (Å) | 129.59 | 130.7 |
| *c* (Å) | 278.48 | 278.9 |
| α=β=γ (°) | 90 | 90 |
| **Data collection** | | |
| X-ray source | APS 23IDD | APS 23IDD |
| Wavelength (Å) | 1.0 | 1.0 |
| Resolution range (Å) | 49.16- 3.35 | 47.75 - 3.10 |
| (Highest res. shell) | (3.51 - 3.35) | (3.22 - 3.10) |
| Completeness (%) | 99.9 (100) | 100(100) |
| *I*/σ(*I*) | 7.2 (0.8) | 10.1 (1.0) |
| CC1/2 | 0.99 (0.55) | 0.99 (0.35) |
| *R*pim | 0.12 (1.8) | 0.07 (0.96) |
| **Refinement statistics** | | |
| *R*-work (%) | 0.24 | 0.23 |
| R-free (%) | 0.28 | 0.26 |
| Clash score | 5.34 | 4.41 |
| Ramachandran plot | | |
| Most favored (%) | 95.0 | 94.4 |
| Additional allowed (%) | 4.4 | 5.2 |
| Disallowed (%) | 0.6 | 0.3 |
| rms bonds (Å) | 0.002 | 0.002 |
| rms angles (deg) | 0.442 | 0.46 |
| PDB code | xxx | xxx |

**Methods:**

**Molecular biology.** The $K_v$1.2-2.1 and the $K_v$beta2.1 genes were a kind gift from Dr. Roderick MacKinnon (The Rockefeller University).(1) $K_v$1.2-2.1 gene was cloned into pAMV vector (kindly provided by Dr. Ming Zhou, Baylor College of Medicine) for electrophysiology experiments and into the pPicZ-C vector (ThermoFisher) for protein expression in *Pischia pastoris*. Site-directed mutagenesis was carried out by PCR based mutagenesis and confirmed through DNA sequencing. Complementary RNA (cRNA) was transcribed using the mMessage mMachine kit (ThermoFisher) and purified using the RNeasy kit (Qiagen).

**Electrophysiology.** *Xenopus laevis* oocytes were provided by Dr. Michael Danilchik (OHSU, protocol # IP00000214), or purchased from Ecocyte Biosciences. Oocytes were injected with 50 nL (50-600 ng) of cRNA and ionic currents were measured 1-4 days after injection using two-electrode voltage clamp (TEVC) on an OC-725 amplifier (Warner). Recordings were carried out either in 100 mM $K^+$ (96 mM KCl, 2 mM NaCl, 5 mM HEPES-KOH, 2 mM $MgCl_2$, pH 7.5) or in 1 mM $K^+$ solution (1 mM KCl, 117 mM NaCl, 0.3 mM $CaCl_2$, 1 mM $MgCl_2$, 5 mM HEPES-NaOH pH 7.5). Glass electrodes used were filled with 3 M KCl and had 1-3 MΩ resistance. Data were sampled at 10 kHz and filtered at 1 kHz. The time constants for inactivation were determined by fitting the decay in current to a single exponential. All data reported were collected from 5-10 oocytes and from at least 3 separate batches of oocytes.

**Protein expression, purification and crystallization.** The $K_v$1.2-2.1-3m and $K_v$1.2-2.1-2m construct in the pPicZ plasmids carried an N- terminal $His_{10}$ tag and a thrombin protease cleavage site. The $K_v$1.2-2.1-3m and $K_v$1.2-2.1-2m pPicZ plasmids were linearized with BglII and ligated with the $K_v$beta 2 gene (36-367 residues on a BglII/BamHI DNA fragment). Plasmids with both the channel and the beta subunit were linearized with PmeI and transformed into *P. pastoris* strain (SMD1163) by electroporation. Transformants were selected on YPDS (yeast extract, peptone, dextrose and sorbitol)(2) plates containing 800 μg/ml of Zeocin. The transformants were grown in liquid culture and tested for expression using anti-His western blotting (Proteintech). Transformants showing good expression were selected and stored as glycerol stocks at -80 $^oC$.

For protein expression, 10 mls of an overnight culture grown in YPD (yeast extract, peptone, dextrose)(2) medium with 200 μg/ ml zeocin was used to inoculate 1 L of BMGY (Yeast nitrogen base, $KPO_4$, pH 6.5 and glycerol)(2) medium with 50 μg/ ml zeocin and grown at 30 $^oC$ for 24 hours. The cells were then pelleted by centrifugation (1500g, 10 min) and transferred to BMMY medium (Yeast nitrogen base, $KPO_4$, pH 6.5 and methanol)(2) with 25 μg/ ml zeocin. After 24 hours, 0.5% (v/v) methanol was added to the culture to induce protein expression and growth was continued for an additional 24 hours. The cells were then pelleted by centrifugation (4500g, 20 min) and frozen in liquid $N_2$ until use.

For purification, the frozen cells were lysed by milling (MM400, Retsch Inc.). Six cycles of milling at 25 Hz for 3 min were carried out. The cells were kept at low temperatures between milling cycles by cooling in liquid nitrogen. The cell powder obtained after milling was solubilized (1gm/ 5ml) in 50 mM Tris-HCl, pH 7.5, 150 mM KCl, 2 mM TCEP (Tris(2-carboxyethyl)-phosphine hydrochloride, 10 mM β-mercaptoethanol (βme), 0.05 mg/ml Deoxyribonuclease I, 1 mM $MgCl_2$, 1 mM PMSF, 1 μg/ml leupeptin, 0.1 μg/ml pepstatin, 1 μg/ml aprotinin and 0.1 mg/ml Soy trypsin inhibitor. The pH of the suspension was adjusted to 7.5 with KOH and the membranes were solubilized with 1.5 % (w/v) n-dodecyl-β-D-maltopyranoside (DDM) for 3h at room temperature. The un-solubilized material was separated by ultracentrifugation (1,00,000g, 50 min). The supernatant following ultracentrifugation was added to cobalt beads (1.5 ml cobalt resin/ 40 ml) pre-equilibrated with column buffer (50 mM Tris-HCl, pH7.5, 150 mM KCl, 2 mM TCEP, 10 mM βme and 5 mM DDM). The bead slurry was overlaid with argon gas and incubated overnight at 4 $^{o}$C with gentle rotation. After incubation, the beads were collected on a column, washed with 20 volumes of column buffer containing 0.1 mg/ml lipids (3:1:1 of POPC (1-palmitoyl-2-oleoyl-sn-glycero-3-phosphocholine): POPE (1-palmitoyl-2-oleoyl-sn-glycero-3-phosphoethanolamine): POPG (1-palmitoyl-2-oleoyl-sn-glycero-3-phosphoglycerol) and 20 mM imidazole and the bound protein was then eluted with 400 mM imidazole. The eluted protein was supplemented with 10 mM DTT, concentrated using a Millipore Amicon Ultra 100 K device and further purified on a Superdex S200 column in 20 mM Tris-HCl, pH 7.5, 150 mM KCl, 2 mM TCEP, 10 mM DTT, 1 mM EDTA, 0.1 mg/ml lipids (3:1:1 of POPC: POPE: POPG), 3 mM Cymal-6 and 3 mM Cymal-7. The fractions containing both the channel and the beta subunits were pooled and concentrated to 15 mg/ml.

For crystallization, 8 mM CHAPS detergent (Anatrace) was added to the protein sample and incubated at room temperature for 45 min. Crystallization was carried out by the hanging drop vapor diffusion method using a crystallization solution of 100 mM Tris-HCl, pH 8.2- 8.8 and 26-36% PEG400 and set up using a Mosquito Crystal system (TTP Labtech). A 1:1 ratio of protein to crystallization solution was used for growing the $K_v$1.2-2.1-3m crystals while a ratio of 2:1 was used for $K_v$1.2-2.1-2m. The rod-shaped crystals generally appeared after 4-5 days. For cryo-protection, the PEG400 concentration in the well solution was increased to 35% (if necessary) and incubated for an additional 24 hours. The crystals were harvested and frozen in liquid $N_2$.

**Solving the $K_v$1.2-2.1-3m crystal structure.** Diffraction data for the $K_v$1.2-2.1-3m crystals were collected at Advanced Photon Source (beamlines 23ID-B and 23ID-D). The crystals were sensitive to radiation damage. Multiple datasets were collected, processed using XDS(3) and analyzed using Pointless(4) and Aimless(5). Sectors from the data sets that contained high resolution diffraction data were merged using BLEND(6) to obtain a complete data set to 3.35 Å. This dataset had a completeness of 100% in the outer-shell (3.51-3.35) with an I/σ of 0.8 and a $CC_{1/2}$ of 0.55 along with minimal anisotropy (3.42 Å in *h,k* and 3.35 Å in *l* planes). The $K_v$1.2-2.1-3m crystallized in the same $P4\,2_1\,2$ space group as the $K_v$1.2-2.1 channel but with slightly smaller unit cell parameters (129.59, 129.59, 278.48).

The $R_{free}$ flag from the dataset for the $K_v$1.2-2.1 structure (pdb: 2r9r)(1) was transferred to the $K_v$1.2-2.1-3m dataset before molecular replacement. For molecular replacement, the $K_v$1.2-2.1 structure with the selectivity filter (residues 370-376), cofactors, lipids and water molecules deleted was used as the search model. Molecular replacement was carried out using Phaser.(7) The asymmetric unit consists of 2 copies of the channel and the beta subunits. The channel subunit consists of the transmembrane TM and the T1 domains. The best molecular replacement solution obtained consisted of the TM domain of 1 channel subunit along with 2 T1 domains and 2 beta subunits. Attempts to place the TM region of the second channel subunit were not successful.

We refined the molecular replacement solution by Jellybody refinement in Refmac(8) followed by multiple cycles of manual structure adjustments using COOT(9) and refinement using Phenix.(10) The electron density in the T1 domains and the beta subunits was very clear and allowed unambiguous placement of the protein chain into the electron density. Similarly, the TM5, pore helix and the TM6 helices were easily modelled into the electron density. The electron density in the selectivity filter residues Y373-D375 was weak and so these residues were not modelled at this stage. The electron density for the voltage sensor domain (VSD) was weak compared to the TM segments in the pore domain but was sufficient for the unambiguous placement of the TM1-TM4 segments in the electron density. The electron density for the loops between the transmembrane helices in the VSD was very poor and so these loops were not modelled. For the Y373-G374-D375 region of the selectivity filter, we calculated Polder(11) omit maps using Phenix, which allowed us to model the amino acid side chains for Y373 and D375. Electron density for the C$\alpha$ and CO of G374 was not observed and so the placement of G374 is speculative. An omit map calculated at this stage showed clear density for the TM regions in the second channel molecule. We placed TM domain from the first channel molecule into this density and refined the structure by jellybody refinement in Refmac followed by manual adjustments in COOT and further refinement in Phenix. The cofactors NAP, lipid behind the selectivity filter and ions/water in the selectivity filter were added. The final model includes residues 36- 361 of the beta subunits (chains A and C). In the channel subunits (Chains B and D), the model includes all the residues in the TM segments while residues in the linker region connecting T1 domain and VSD (133- 142 in B and 134- 147 in D), the TM1-TM2 loop (192-217 in B and 191-218 in D) and the TM3-TM4 loop (273-286 in chain B and chain D) were not modelled. Figures were prepared using Pymol(12) or Chimera.(13) The solvent accessible area (SASA) for $K_v$1.2-2.1 and $K_v$1.2-2.1-3m were calculated using Qt-PISA program in CCP4.(14)
